# Detection of clade 2.3.4.4b H5N1 high pathogenicity avian influenza virus in a sheep in Great Britain, 2025

**DOI:** 10.1101/2025.06.27.661969

**Authors:** Ashley C. Banyard, Holly Coombes, Jacob Terrey, Natalie McGinn, James Seekings, Benjamin Clifton, Benjamin C. Mollett, Cecilia Di Genova, Pia Sainz-Dominguez, Laura Worsley, Raquel Jorquera, Elizabeth Billington, Edward Fullick, Audra-Lynne Schlachter, David Jorge, Alejandro Núñez, Marco Falchieri, Joe James, Scott M. Reid

## Abstract

Clade 2.3.4.4b H5N1 high pathogenicity avian influenza virus (HPAIV) continues to pose a significant global threat, affecting wild and domestic avian populations, and mammalian species. In early 2024, H5N1 HPAIV was detected in dairy cattle in the United States of America, where it has continued to circulate, with sporadic detections also reported in other ruminant species. The detection of high viral loads in milk from infected cattle, has resulted in several human infections, underscoring the zoonotic potential of these viruses. In response, several countries have intensified surveillance in non-avian species to evaluate the potential for undetected viral circulation in captive mammals. In Great Britain, bulk milk tank testing of cattle and targeted surveillance of captive mammalian species on an infected premises is undertaken in accordance with the outcome of a rapid risk assessment undertaken to determine the epidemiological links between the poultry and captive mammals. A result of this testing was the first recorded detection of clade 2.3.4.4b H5N1 HPAIV in a sheep in March 2025, identified on an infected poultry premises in Great Britain. An initial seropositive result in a single ewe triggered further investigation, confirming serological positivity across repeated sampling and the presence of viral RNA in milk samples. This detection was confined to a single animal and was likely attributable to proximity to infected poultry and a presumed heavily contaminated environment. The implications of this ruminant detection are discussed in the context of interspecies transmission and surveillance strategies.

## Introduction

Clade 2.3.4.4b H5N1 high pathogenicity avian influenza virus (HPAIV) continues to cause significant mortalities in both birds and mammals globally [1, 2] with outbreaks decimating wild bird populations, the poultry sector, and populations of terrestrial and marine mammals [2,3]. High wild bird mortality rates have also affected species of conservation concern [2,4] increased environmental pressure leading to the infection of terrestrial and aquatic mammals [3]. Infection of captive mammals has also been reported in zoological collections, household settings and farming sectors and have included captive bush dogs [5], farmed mink and foxes [6,7], household cats [8], and most notably farmed dairy cattle in the United States of America (USA) [9,10].

From a ruminant perspective, H5N1 HPAIV (genotype B3.13) was first reported in dairy cattle in the USA on 25^th^ March 2024 [11] although phylogenetic analyses have proposed initial infection of cattle in late 2023. Since this detection, B3.13 has spread in cattle, forming a discrete lineage, yet becoming extinct in wild birds [12]. The infection of dairy cattle has significantly elevated the zoonotic risk of these viruses, due to; (**i**) the detection of high viral loads in milk from infected cattle, and (**ii**) concerns around mammalian adaptations that risk increasing viral fitness in humans [10, 13, 14]. A second H5N1 HPAIV genotype (D1.1-detected in wild birds across the USA [17]) was also reported in cattle in the state of Nevada, USA on the 31^st^ January 2025 [15], and separately in the state of Arizona, USA on 13^th^ February 2025 [16]. While the spillover dynamics of HPAIV into cattle remains unclear, these detections clearly demonstrate infection risk. From a zoonotic perspective, of the 70 cases of human infection reported in the USA with H5N1, 41 have been associated with exposure to infectious material in a dairy setting [11]. Alongside dairy cattle, H5N1 has also been sporadically reported in other farmed mammals including alpacas and goats [12, 18], as well as captive mammals present on farms such as cats [19].

Alongside infection of dairy cattle with genotypes B3.13 and D1.1 [18, 20], experimental studies have demonstrated that cattle infection is not constrained by genotype with American and European strains being able to induce similar infection characteristics in cattle and other mammalian species [21]. These findings have led to enhanced surveillance for HPAIV in mammals, particularly where there are linkages to infectious material on infected poultry premises. However, to date, H5N1 HPAIV has not been detected in cattle outside of the USA [22–25]. In Great Britain (GB), national bulk milk tank sampling of a representative proportion of dairy cattle herds was undertaken. Over 500 bulk-milk samples, from 455 farms across GB, were all negative for HPAIV [26]. In addition, targeted surveillance is being undertaken for captive mammals co-located with infected poultry where the risk from HPAIV is considered greater than the baseline risk posed by wild birds. Within GB, the detection of influenza of avian origin in mammalian species has been limited to sporadic infections, including foxes, otters and sporadic detections in marine mammals [27]. There has only been a single detection of H5N1 in captive mammals in GB in a single group of captive bush dogs [5, 28]. However, as the clade 2.3.4.4b H5N1 panzootic continues to impact both wild and captive avian species, the risk to mammals co-located with infected poultry remains.

Here, the detection of H5N1 in a backyard flock including chickens, ducks, turkeys and geese is described whereby virus detection in avian species led to risk-based sampling of mammals and the detection of evidence for clade 2.3.4.4b H5N1 HPAIV in a single sheep. Factors leading to this detection are described.

## Materials and methods

### Clinical investigation, post-mortem (PM) examination, tissue sampling and histopathological analysis

Samples from birds were collected as described previously [29]. Oral and rectal swabs samples were collected from a proportion of sheep alongside blood sampling. Later sampling included milk from individual animals. Following molecular and serological detection of viral material in a ewe, a full PM examination and tissue sampling was undertaken. Tissue samples were processed molecular and histopathological assessment. To homogenise tissues, approximately 0.5 grams of each tissue was processed into Precellys Hard tissue Homogenizing Tubes (CK28; Bertin Technologies, France) with 0.5ml of phosphate-buffered saline (PBS). Samples were homogenised for 2 min at 7000 RPM using Precellys24 homogeniser (Bertin Technologies, France). Supernatants were centrifuged at 10,000g for 30 seconds, and clarified supernatants were processed for extraction of viral RNA (vRNA) for molecular analyses [30].

For histopathology samples were processed as described previously [31]. Twenty-two individual wax-blocks were prepared from mammary gland (11 for each gland) to assess for infection. Haematoxylin and eosin (H&E) and immunohistochemistry (IHC) was undertaken as described previously [31] and stating assessed using a previously established criteria [32]. Specificity of immunolabelling was assessed using positive control sections and by replacing the primary antibody with a matching mouse IgG isotype in test sections. Additionally, Gram Twort staining was performed in mammary gland sections.

### Virological investigation

#### RNA extraction and Molecular Analysis

RNA was extracted from samples as previously described [30]. Extracted RNA was assessed for vRNA using RT-PCR assays including those targeting matrix (M) [33], a HPAIV H5 [34] and/or NA [35, 36]. RT-PCR Cq values < 36.00 were considered as positive. Samples with Cq ≥36 were considered negative. A standard curve was generated using a ten-fold dilution series of titrated H5N1-21/22 HPAIV RNA as previously described [30]. RNA was also extracted from a range of clarified tissue homogenates as described earlier.

#### Attempted virus isolation (VI)

For VI, 100 µl of the sample material was diluted with 100 µl of PBS and inoculated into the allantoic cavity of three specific pathogen-free (SPF) embryonated chicken eggs (ECEs)[37, 38]. The allantoic fluid from one ECE was tested for the presence of virus at 2 days post inoculation (dpi) using the haemagglutination assay as previously described [37]. Fluids exhibiting haemagglutination activity ≥16 were considered positive, whereas those with activity <16 were classified as negative. Allantoic fluids testing negative at 2 dpi were subjected to a second passage by inoculation into additional ECEs. At six dpi, allantoic fluid from all ECEs was harvested and tested by HA. Samples yielding negative haemagglutination activity following this protocol were deemed negative for infectious virus. Attempts were also made to concentrate virus present in milk by the addition of chicken red blood cells (cRBCs) [39].

#### Serological assessment

Sera was separated from blood samples as described previously [40]. Hemagglutination inhibition (HI) assays were undertaken using A/Chicken/Wales/053969/2021 (H5N1) (clade 2.3.4.4b H5N1) antigen [37]. In addition, batches of serum which included an HI positive sample were run on two commercial ELISA tests were used as per manufacturer’s instructions (ID Screen^®^ Influenza A Antibody Competition Multi-species ELISA and the ID Screen^®^ Influenza H5 Antibody Competition 3.0 Multi-species ELISA (Innovative diagnostics).

#### Genomic analysis

Viral genome sequences were generated from the composite milk sample using Oxford Nanopore Technology, by adapting a method previously described [41, 42]. For all avian samples extracted vRNA was processed using the SuperScript III One-Step RT-PCR kit (Thermo Fisher Scientific). Extracted RNA from the milk samples were converted to double stranded cDNA and amplified in triplicate with all three aliquots of amplified cDNA being pooled and concentrated by ethanol precipitation.

To complete sequences for the milk PA, PB1 and HA segments, segments were converted to cDNA and amplified individually using segment specific primers (Supplementary Table 1). Amplified cDNA was purified with Agencourt AMPure XP beads (Beckman Coulter) prior to sequencing library preparation using the Native Barcoding Kit (Oxford Nanopore Technologies) and sequenced using a GridION Mk1 (ONT), according to manufacturer’s instructions. Assembly of the influenza A viral genomes was undertaken as described previously [41]. Raw reads from the milk, PB1, PA and HA samples were combined prior to assembly. Comparison of the study-derived sequences and contemporary H5 sequences was undertaken against all avian H5 sequences available on GISAID between 1^st^ January 2020 and 23^rd^ April 2025. All sequences were aligned on a per segment basis using Mafft v7.520 [43] and trimmed against a reference using SeqKit v2.5.1 [4]. The trimmed alignments were used to a infer maximum-likelihood phylogenetic tree IQ-Tree version 2.2.3 [44] along with ModelFinder [41] and 1,000 ultrafast bootstraps [43]. Inference of molecular-clock phylogenies were performed using TreeTime v0.10.1 [45] and trees were visualised using R with ggplot2 [46] and ggtree version 3.14.0 [47]. Sequences were genotyped from phylogenetic trees by comparison to known reference sequences for all genotypes currently circulating in GB.

All sequences were assessed for the presence of adaptive mutations by aligning on a per segment basis using MAFFT v7.520 [43] and manually trimming to the open reading frame using Aliview version 1.28 [48]. Trimmed sequences were translated to amino acids and visually inspected for mutations. All influenza sequences generated in this study are available through the GISAID EpiFlu Database (https://www.gisaid.org, Supplementary Table 2).

## Results

On 18^th^ February 2025 HPAI H5N1 was confirmed on a backyard mixed poultry holding (n=60 poultry) in Yorkshire. The infected premises consisted of 34 chickens (*Gallus gallus domesticus*), 5 turkeys (*Meleagris gallopavo*), 19 ducks (*Anas platyrhynchos domesticus*) and 2 geese (*Anser anser domesticus*) (Figure 1A & B). The birds had been housed since the housing order came into effect on 23^rd^ December 2024 as the infected premises lies within the Avian Influenza Prevention Zone (AIPZ) [49]. The poultry were housed in four groups; Group A comprised 31 chickens in a stable, Group B comprised 19 ducks and two geese in another stable, Group C comprised five turkeys in another shed, and Group D comprised 3 chickens in a wooden coop in the garden. Additionally, the premises had one household cat (*Felis catus*) and 26 sheep (*Ovis aries*) (16 breeding ewes, 9 lambs and 1 ram). At the time of the report case (Figure 1A), one group of sheep were housed next to the turkey shed (Group E), another group (which included the affected ewe) were housed next to the duck and goose shed (Group H) with the remainder in the field adjacent to the garden (Group I). There were no linked premises nor commercial activities reported for this premises. Eggs and meat from the premises were only consumed by the keeper within the window of suspected infection.

**Figure 1.**
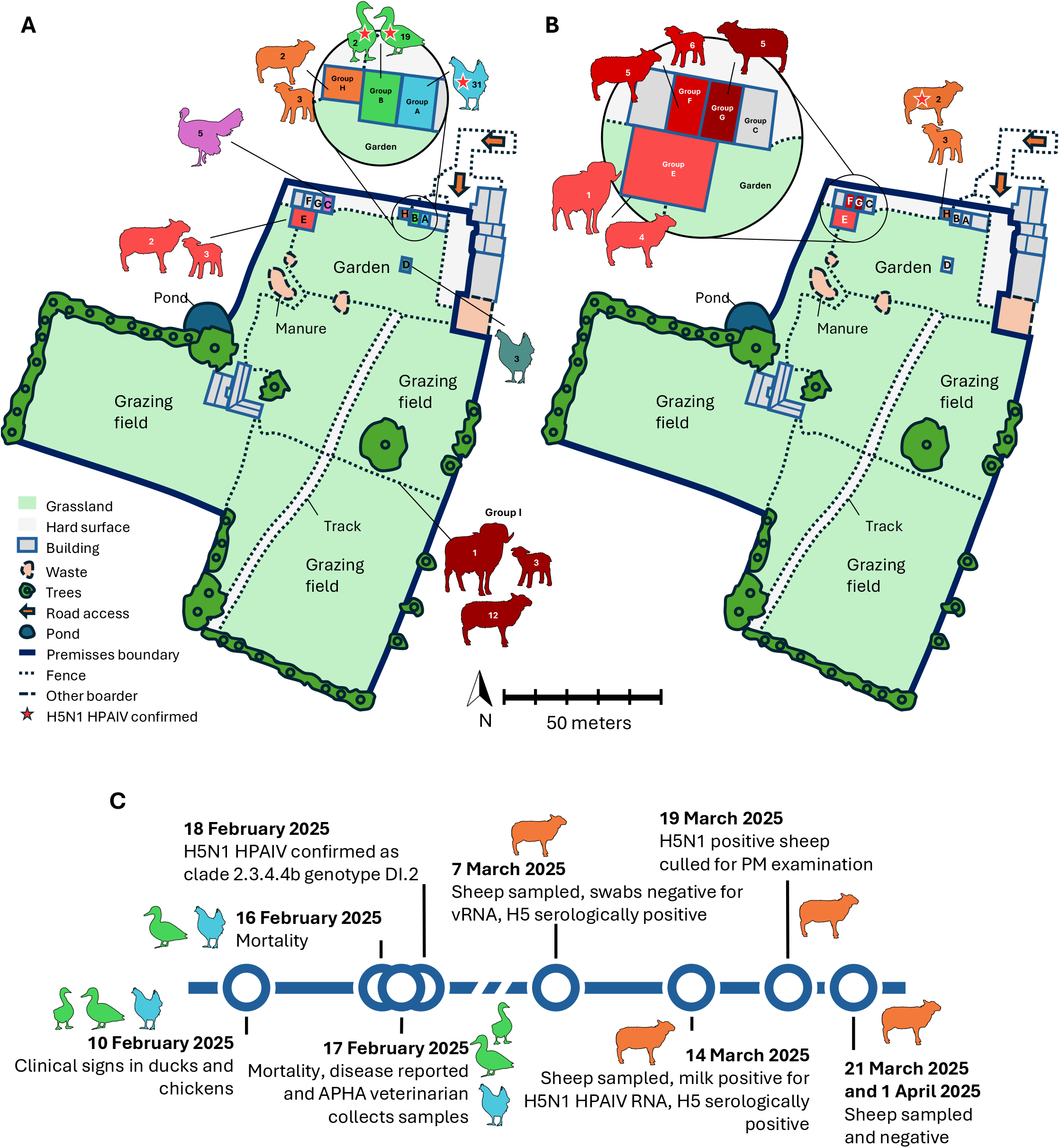
Map of the infected poultry premises and timeline of key events. The infected poultry premises indicating geographical features and animal locations and numbers during veterinary visits **(A)** at the time of poultry sampling (17th February 2025), **(B)** at the time of initial sheep sampling (7th March 2025). **(C)** A timeline of key events during the event.

Although the birds were housed indoors, overall biosecurity at the premises was assessed as poor. Structural deficiencies such as gaps in roofs, doors, and windows were identified, and no disinfectant footbaths were in use either at the entrance to the premises or between different sheds. The poultry sheds were not secure against wildlife ingress; entry points such as damaged windows, holes in the roof, and visible gaps in gates allowed potential access. Evidence of rat activity was also present, and the keeper reported observing a rat feeding on the head of a deceased chicken, although no samples were available from rats on site. The birds were bedded daily, and the interiors of the poultry sheds were clean and dry. Mains-supplied drinkers were available within each shed. Although wild bird activity on site was reported to be low, sightings of wild birds flying overhead had been noted and there were bird feeders in the garden area near the sheds housing the poultry and sheep. Following the onset of avian mortality, recently deceased birds were stored in a garden waste bin.

The infection event in avian species began with initial signs of inappetence observed on 10^th^ February 2025 in both the ducks and chickens, with a decrease in egg production among the chickens (Figure 1C). By 16^th^ February 2025, one chicken and one duck had died, and one of the two geese was exhibiting neurological signs consistent with clinical disease, including head tremors. On 17^th^ February 2025, the clinically affected goose was found dead, and a further two chickens and one duck had succumbed to the infection. The owner reported suspicion of disease on 17^th^ February 2025 (Figure 1C). An official veterinarian from the Animal and Plant Health Agency (APHA) attended the premises the same day. Upon inspection, two of the four bird groups (Groups A and B) were found to be affected (Figure 1A). Two ducks displayed clinical signs suggestive of a notifiable avian disease (NAD); one exhibited “snicking” (a respiratory sign analogous to sneezing in humans), while the other appeared lethargic with ocular discharge. Additionally, one chicken was noted to be lethargic. A PM examination of one chicken revealed no significant gross lesions. However, samples were collected including oropharyngeal and cloacal swabs from 8 chickens (Group A), with 2 whole heads submitted for brain sampling; oropharyngeal and cloacal swabs from 11 ducks, along with 2 blood samples (from duck 18 and 19); and oropharyngeal and cloacal swabs from 1 goose (Group B). One of the clinically affected ducks died later on 17^th^ February 2025, and by the following day, the remaining clinically affected duck and the affected chicken had also died (Figure 1C).

Swab samples tested positive by RT-PCR for the influenza A virus (M-gene), H5 HPAIV (HP-H5), and N1 vRNA in 50% (4/8) of the chickens, 100% (11/11) of the ducks, and the single goose (1/1) (Supplementary Table 3). Serological testing of duck blood samples by HAI revealed one (duck 18) out of two samples to be positive for H5 antibodies against clade 2.3.4.4b H5N1 antigen, with an HI titre of 1/64. These laboratory findings, in conjunction with the observed clinical signs, led to the official confirmation of H5N1 HPAI in avian species on the premises on 18^th^ February 2025 (Figure 1C). Subsequent genetic assessment identified all poultry genomes as a clade 2.3.4.4b H5N1 HPAIV, belonging to the DI.2 genotype, in accordance with the established genotype classification system [50].

An epidemiological assessment was undertaken for risk factors for mammalian infection that resulted in sampling of the sheep for diagnostic evaluation (Figure 1B). On the 7^th^ of March, ten of the 26 sheep were sampled; the remaining animals were not sampled due to welfare concerns, as they were heavily pregnant. Nasal and rectal swabs collected from the sampled sheep (n = 10) (across Group H, G and F; Figure 1B) tested negative for vRNA by RT-PCR using the M-gene, H5-HP and N1 assays. Notably, serological testing of blood samples revealed that one ewe in Group H (Figure 1B) had detectable H5-specific antibodies, with a HI titre of 1:80 (Supplementary Table 4). Unaccredited ELISAs also returned positive results from this serum, indicating the presence of antibodies reactive to NP and H5 antigens. This initial positive serological detection led to repeat sampling of that sheep which was undertaken on the 14^th^March 2025 and although the animal was negative for vRNA from nasal and rectal swabs (Supplementary Table 5), the sheep was again positive for H5 reactive antibodies by HAIT (having a titre of 1/160), and the serum was positive by both ELISA tests. A composite milk sample (collected from both halves of the udder) tested positive for H5N1 HPAIV vRNA by M gene, HPH5 and N1 PCR assays. The milk sample was also assessed by competitive ELISA for NP and H5 and was positive in both tests.

Following the detection of H5-specific antibodies and H5N1 HPAIV vRNA in the milk, the affected ewe was culled on the 19^th^ of March 2025, and a full PM examination was undertaken. All tissue samples (n = 28) and swab samples (n = 12) tested negative for HPAIV RNA by M gene, H5-HP, and N1 RT-PCRs. However, a further milk sample taken from the ewe at cull tested positive for H5N1 vRNA across all three RT-PCR assays, consistent with previous findings (Supplementary Table 5). Subsequently, tissue samples were further homogenised and reassessed for vRNA. Borderline detection of vRNA was observed in the homogenised right mammary gland tissue, but this result could not be consistently reproduced upon repeat RNA extraction and testing. Histological examination was conducted on these tissues but no lesions consistent with active mastitis or viral replication within the glandular tissue were observed, nor were there any definitive histological features indicative of prior viral exposure. Histopathological examination revealed areas of multifocal lymphoplasmacytic infiltration around alveoli and ducts in one half. In contrast, multifocal regions of the other half exhibited acute neutrophilic alveolar exudation, but not showing intraluminal epithelial sloughing and cellular debris in mammary alveoli described in active infection in cattle. IHC for the presence of viral antigen was also undertaken on mammary tissue sections; however, all sections were negative for viral antigen. Gram staining (Twort counterstain) of mammary gland samples did not reveal the presence of bacterial colonies.

VI was attempted on both H5N1 HPAIV positive milk samples using ECEs. However, infectious virus could not be isolated. We also attempted to sequester and concentrate potential virus in the milk samples using cRBCs. However, no infectious virus could be isolated following this treatment.

Full viral sequences were obtained from the milk sample for all segments except the PB1, which contained a large gap between nucleotides 333 and 786, as well as some smaller gaps between 190 and 330. The available sequence was compared to those derived from the infected birds and identified as a genotype DI.2 H5N1 clade 2.3.4.4b virus (Supplementary Table 2), clustering with other DI.2 isolates detected in GB during the same period (Supplementary Figure 1). Further analysis of positive material from all infected birds (n = 21) on the premises revealed >99.9% sequence identity across all segments. The viral sequence from the sheep contained nine amino acid substitutions relative to the avian viral sequences (Supplementary Table 6). Two changes were identified in the HA protein (D171N and D277G); neither is associated with increased zoonotic potential, although D171N has been previously reported in genotype B3.13 H5N1 sequences from cattle in the USA and may represent a ruminant-associated adaptation (Figure 2) [12, 51]. Additional substitutions distinguishing the sheep and avian sequences included PB2 N456D, NA L75F and V114M, PA L335F and PB1 M290V, K577E, and Q688H. Notably, previously reported cattle-adaptive mutations, such as PB2 E362G, M631L, and E677G; PB1 N642S; and PA A448S [12, 52], were absent from the sheep-derived sequence. No mutations were detected in either the NP, MP or NS genes when compared with the avian reference sequence (Figure 2).

**Figure 2:**
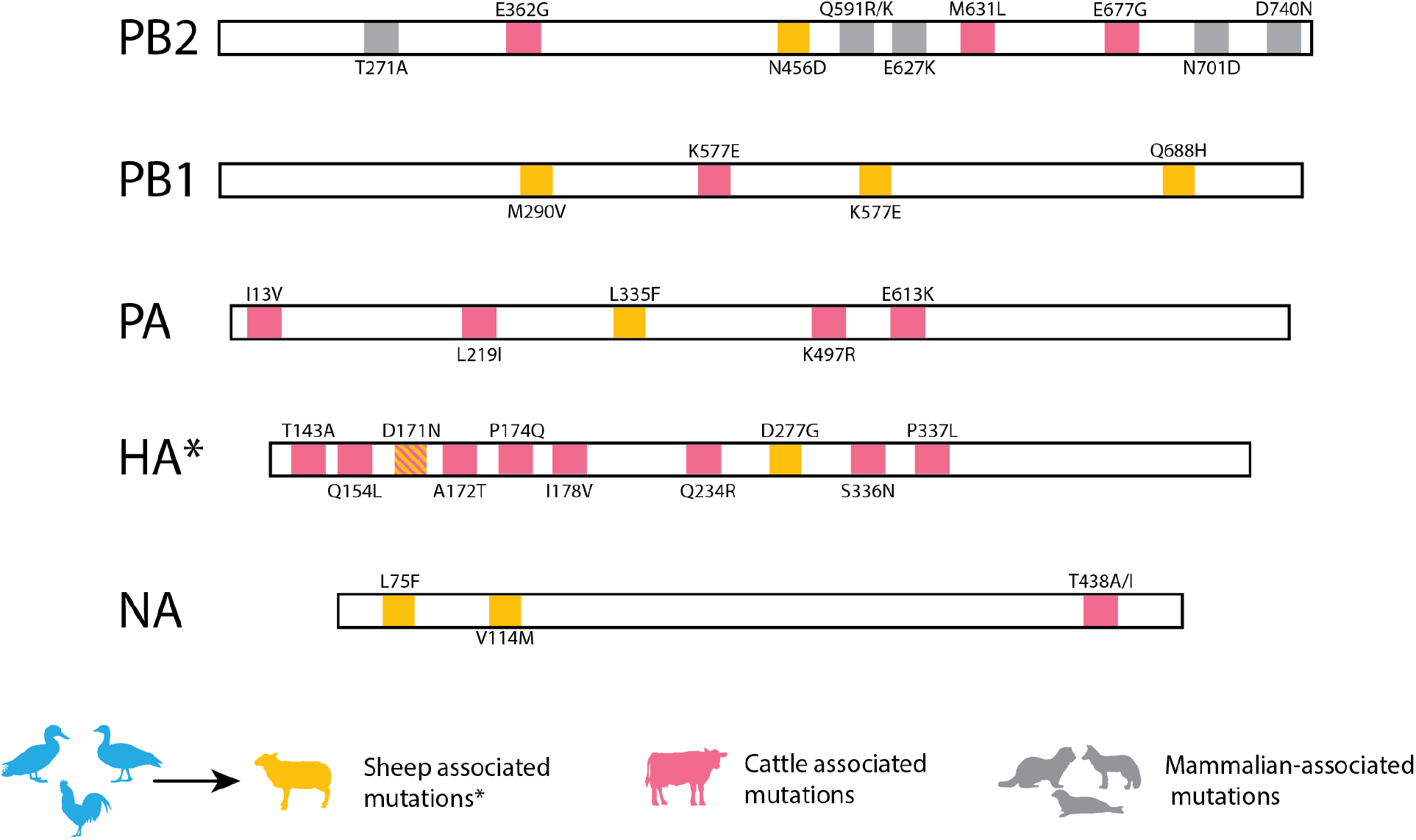
Genetic changes in sheep sequences. Amino acid mutations in A/Sheep/England/023754/2025 consensus sequence compared to co-located avian species. Cattle mutations from [12, 51, 52]. NP, MP and NS segments shared 100% identity.

## Discussion

The detection of clade 2.3.4.4b H5N1 vRNA in a sheep represents the first reported case of infection with this virus in this species globally, and the first detection in a ruminant outside of the USA. Previous detections of H5N1 in mammalian species have been attributed to elevated environmental infection pressure. Transmission pathways are often apparent, particularly among scavenging species, where this mammal infection is presumed to be the result of predation on infected carcasses [2, 3].

However, this unusual detection in a sheep raises the possibility of alternative routes of infection. The ewe was co-located on an infected holding where chickens, ducks, and geese had tested positive for H5N1, and had been housed since December 2024 in a shed directly adjacent to the one housing both the ducks and geese (Group B). Importantly, the presence of infection in *Anseriformes* species (ducks and geese) increases the likelihood of a heavily contaminated environment, given their known capacity for extensive environmental viral shedding [30, 53, 54]. Experimental work consistently demonstrates that infected ducks release high viral loads into the environment, contaminating water and surfaces extensively. Based on this evidence, the lack of mitigating biosecurity measures in place between the poultry and the sheep managed by a single keeper who oversaw their daily management, and the common DI.2 sequence results isolated from the poultry and the ewe, it is reasonable to infer that the ewe may have acquired infection through indirect contact with the infected poultry shedding large amounts of HPAIV into the environment. Other hypotheses, such as direct contact with infected wild birds or vermin, are less likely. Indirect contact with wildlife was identified as the most likely source of infection for the poultry on this premises.

Despite these findings, the precise route of infection in the ewe remains undetermined. However, several potential pathways are plausible, including (**i**) infection though the respiratory or oral routes via contaminated fomites, (**ii**) intramammary infection via contaminated material introduced into the teat by suckling lambs, or (**iii**) direct inoculation into the mammary gland as a result of the ewe lying in virus-contaminated material. Although none of these scenarios can be definitively ruled out, respiratory infection followed by systemic viral dissemination resulting in vRNA detection in milk appears unlikely. Analysis of airborne virus present in commercial poultry sectors has demonstrated relatively low levels of virus present even within densely packed houses were assessed [55]. While the infectious dose for sheep is unknown, considering the density of infected poultry present, the level of virus present in the air was likely low. In addition, if this had occurred, one would expect to observe evidence of systemic infection, such as seropositivity in other sheep or lambs; however, all animals tested were seronegative. Of note, albeit anecdotal, the ewe with detectable vRNA in milk had a reported episode of mastitis on the 3^rd^ March 2025. No additional clinical signs or behavioural abnormalities were reported in the sheep following detection of infection in poultry. At the time of culling on the 19^th^ March 2025, the ewe remained lactating, and milk was again expressed from both teats. No abnormalities in milk consistency or presence of clots were observed. Gross PM examination noted the left mammary gland to be slightly firm and mildly reddened; however, these findings were not considered clinically significant. Milk was collected post-euthanasia and post-swabbing, with samples from both halves of the udder pooled into a single container. Notably, this milk sample tested positive for both anti-NP and anti-H5 antibodies, indicating that the ewe was likely sampled during a phase of viral clearance. This serological profile may explain the failure to isolate infectious virus from the collected samples.

For the second hypothesis (transmission via suckling lambs) to be plausible, one would expect some evidence of infection in the lambs, either through direct mucosal contact with infectious material or ingestion of contaminated milk. However, all lambs tested negative by both swab and serological assays, making this route of transmission less likely. Therefore, the third hypothesis appears most consistent with the available data. Experimental studies have shown that intramammary inoculation with both North American and European genotypes of H5N1 can lead to infection in cattle and is an efficient route of infection, albeit one that generally results in infection restricted to the udder [21]. In such cases, infection is frequently restricted to a single mammary quarter, without resulting in systemic dissemination. Unfortunately, in this case, milk samples were pooled from both teats, precluding any assessment of whether the infection was localised to one mammary gland. This limitation hinders the ability to definitively determine the anatomical extent of infection.

The histopathological findings were unable to provide clear evidence of active viral infection although in one half of the mammary gland subacute to chronic changes may have been consistent with a prior viral infection, the neutrophilic alveolar exudation in the other half would suggest a more recent insult or recent clearance, as no epithelial sloughing and cellular debris in the mammary alveoli associated with active infection [10] were seen. Although extensive sampling and examination of the mammary gland was conducted, some areas where virus was replicating may have been present elsewhere. Further data limitations arose due to logistical constraints associated with sampling pregnant animals. The detection of seropositivity in a single ewe highlighted the need for broader investigation within the flock. However, due to the flock’s lambing status, comprehensive sampling of the entire flock could not be completed and only the initially seropositive ewe was subjected to further diagnostic evaluation at the time. All of the unaffected sheep in the flock were sampled at least once, and all with negative results, with testing completing when the four ewes who did not lamb until late March were found to be negative on 1^st^ April 2025.

The viral sequence obtained from the ewe provides the only definitive evidence that replication occurred within this host as several amino acid substitutions were detected across multiple genes. These changes were absent in viral sequences derived from infected birds on the same premises, which exhibited a high degree of genetic identity among themselves. Such changes strongly support that the virus underwent replication and genetic adaptation within the ewe, consistent with host-specific evolutionary pressure. However, the functional significance of these mutations, particularly regarding viral fitness, host range, or transmissibility, remain unknown.

In conclusion, this study has demonstrated possible infection routes for mammalian species co-located on infected poultry premises. A recent report found that blood samples collected in 2024 from 220 sheep, which had grazed in areas affected by dead and diseased H5N1-infected pheasants in Norway during 2023, contained antibodies against H5 AIV [56]. Interestingly no clinical disease was reported in these sheep [56]. Clearly, such spillover events may be more common than previously recognised and enhanced surveillance is required to demonstrate impact. Detections like this underscore the importance of rapid assessment and dissemination of this information across the global scientific community.

## Acknowledgements

APHA authors were funded by the Department for Environment, Food and Rural Affairs (Defra, UK) and the Devolved Administrations of Scotland and Wales, through the following programmes: SV3400, SV3032, SE2227 and SE2230. This work was also supported by the Biotechnology and Biological Sciences Research Council (BBSRC) and Department for Environment, Food and Rural Affairs (Defra, UK) research initiative ‘FluTrailMap’ [grant number BB/Y007271/1]. Funded by the European Union under grant agreement (101084171) - (Kappa-Flu). Views and opinions expressed are however those of the author(s) only and do not necessarily reflect those of the European Union or REA. Neither the European Union nor the granting authority can be held responsible for them.

