## Supplementary Figures and Tables for "Detection of clade 2.3.4.4b H5N1 high pathogenicity avian influenza virus in a sheep in Great Britain, 2025"

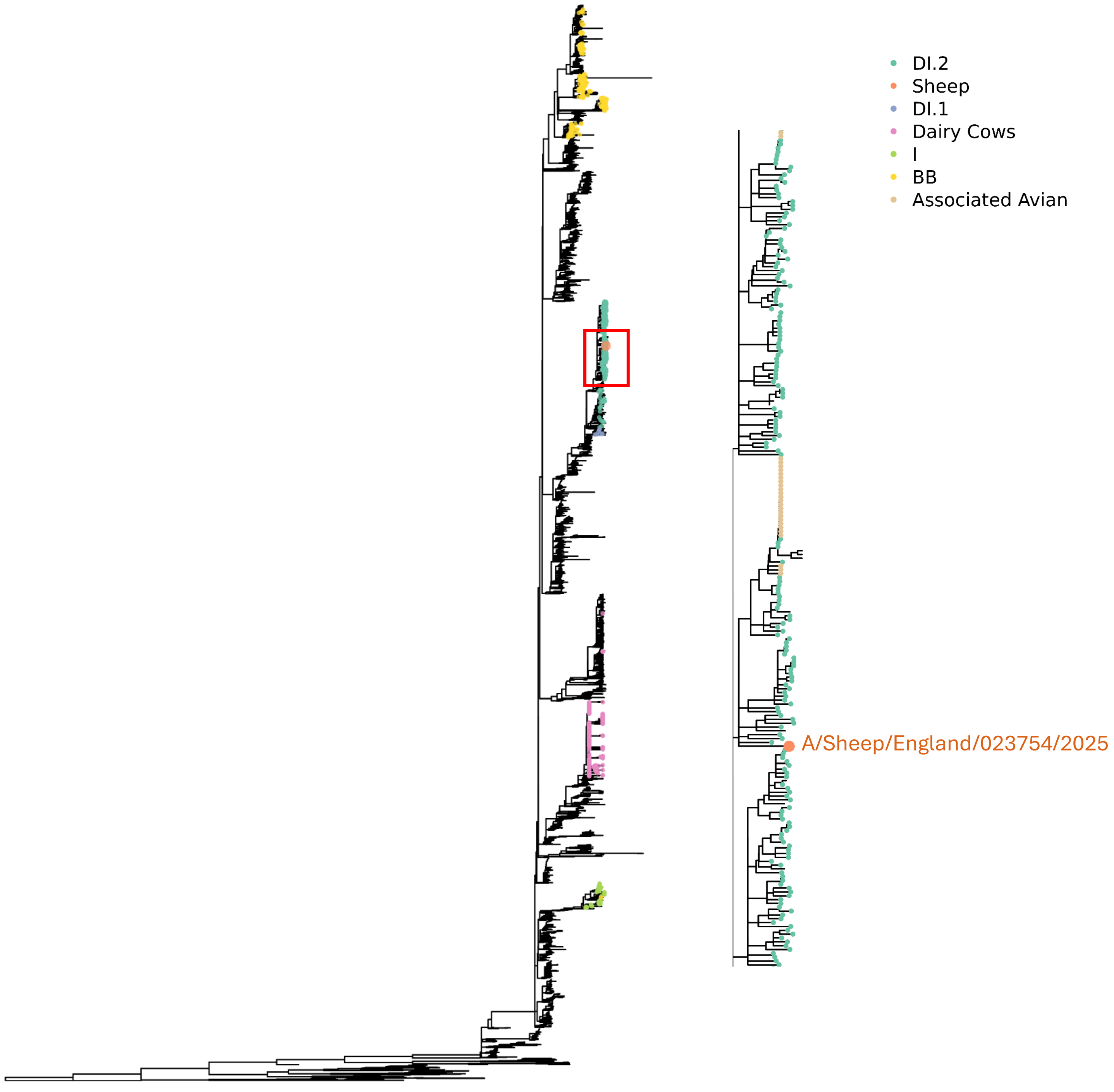


**Supplementary Figure 1: Phylogenetic tree based on nucleotide sequence of HA gene.** Genotypes currently circulating in the UK highlighted along with sequences originating from cattle. Zoomed section highlighting sheep and associated poultry sequences.

**Supplementary Table 1:** Oligonucleotide primers used for cDNA synthesis and amplification for whole genome sequencing.

| Name | Sequence (5' to 3') | Target Segment/gene | Reference |
|---|---|---|---|
| Optil – F1 | TTACGCGCCAGCAAAAGCAG | All | [12]; [13] |
| Optil – F2 | GTTACGCGCCAGCGAAAGCAGG | All | |
| Optil – R1 | GTTACGCGCCAGTAGAAACAAG | All | |
| PB1 - F1 | CWCARATACCWGCAGARATGCT | PB1 | |
| PB1 – F403 | CAGACCTATGACTGGACATTGAA | PB1 | |
| PA-F1 | AGC AAA AGC AGG TAC TGA TCC AA | PA | |
| PA-R1493 | CTG CAT TTG CTT ATC ATT GGA ATC | PA | |
| PA-F982 | GCC CAA CAT CGT GAA ACC ACA | PA | |
| PA-R2233 | AGT AGA AAC AAG GTA CTT TTT TGG A | PA | |
| S1H5N8 | AGCAGGGGTTCACTCTGTCA | HA1 | |
| AS3 | GCTGGATGGCTCCTCGG | HA1 | |
| J4 | GATTTCAACGACTATGAAGAACTG | HA1 | |
| AS1 | AATGATGCGGCAGAGCAGAC | HA1 | |
| KHA1 | CATCAATTTTGAGAGTAATG | HA1 | |
| B2a | TTTTGTCAATGATTGAGTTGACCTTATTGG | HA1 | |
| AS4r | CCCCACAGTACCAAAAGATC | HA1 | |
| AS2r | CCGAGGAGCCATCCAGC | HA1 | |
| J3 | GATAAATTCTAGCATGCCATTCC | HA1 | |
| J3 | GATAAATTCTAGCATGCCATTCC | HA2 | |
| H5-6 | GGGTACCACCATAGCAACGAGCAGGG | HA2 | |
| AS6 | TACCCGCAGTATTCAGAAG | HA2 | |
| J1c | AGTAGAAACAAGGGTGTT | HA2 | |
| J2c | GGTGTTTTTAACTACAATCTGG | HA2 | |
| TK-H5-2r | GGGAGCTCGCCACTGTTG | HA2 | |
| B2a | TTTTGTCAATGATTGAGTTGACCTTATTGG | HA2 | |
| KHA3 | TACCAACCGTCTACCATKCCYTG | HA2 | |

**Supplementary Table 2:** Sequences generated in this study.

| Species | Sample* | Collection Date | Subtype | Virus Strain Name |
|---|---|---|---|---|
| Sheep | Milk | 2025-03-19 | H5N1 | A/sheep/England/023754/2025 |
| Domestic Duck | Op swab | 2025-02-17 | H5N1 | A/domestic_duck/England/014229/2025 |
| Domestic Duck | C swab | 2025-02-17 | H5N1 | A/domestic_duck/England/014230/2025 |
| Domestic Duck | Op swab | 2025-02-17 | H5N1 | A/domestic_duck/England/014237/2025 |
| Domestic Duck | Op swab | 2025-02-17 | H5N1 | A/domestic_duck/England/014241/2025 |
| Domestic Duck | C swab | 2025-02-17 | H5N1 | A/domestic_duck/England/014242/2025 |
| Domestic Duck | Op swab | 2025-02-17 | H5N1 | A/domestic_duck/England/014243/2025 |
| Domestic Duck | Op swab | 2025-02-17 | H5N1 | A/domestic_duck/England/014245/2025 |
| Domestic Duck | Op swab | 2025-02-17 | H5N1 | A/domestic_duck/England/014247/2025 |
| Domestic Duck | C swab | 2025-02-17 | H5N1 | A/domestic_duck/England/014248/2025 |
| Domestic Duck | C swab | 2025-02-17 | H5N1 | A/domestic_duck/England/014270/2025 |
| Domestic Duck | Op swab | 2025-02-17 | H5N1 | A/domestic_duck/England/014232/2025 |
| Domestic Duck | Op swab | 2025-02-17 | H5N1 | A/domestic_duck/England/014235/2025 |
| Chicken | Op swab | 2025-02-17 | H5N1 | A/chicken/England/014249/2025 |
| Chicken | C swab | 2025-02-17 | H5N1 | A/chicken/England/014250/2025 |
| Chicken | Op swab | 2025-02-17 | H5N1 | A/chicken/England/014259/2025 |
| Chicken | C swab | 2025-02-17 | H5N1 | A/chicken/England/014263/2025 |
| Chicken | Op swab | 2025-02-17 | H5N1 | A/chicken/England/014265/2025 |
| Goose | C swab | 2025-02-17 | H5N1 | A/goose/England/014272/2025 |
| Goose | Op swab | 2025-02-17 | H5N1 | A/goose/England/014271/2025 |
| Chicken | Brain | 2025-02-17 | H5N1 | A/chicken/England/014363/2025 |
| Chicken | Op swab | 2025-02-17 | H5N1 | A/chicken/England/014262/2025 |

* C = Cloacal; Op = Oropharyngeal

**Supplementary Table 3**: RT-PCR testing results for H5 HPAIV from captive chickens (Group A), ducks and geese (Group B) sampled on the 17 February 2025 where the sheep were co-located.

| Group | Species | Bird ID | Sample Type | Interpretation from RT-PCR assay | | | |
|---|---|---|---|---|---|---|---|
| | | | | M-gene | H5-HP | N1 | L-gene |
| Group A | Chicken | 10 | Op | Positive | Positive | Positive | Negative |
| | | | C | Positive | Positive | Positive | Negative |
| | Chicken | 11 | Op | Negative | Negative | Negative | Negative |
| | | | C | Negative | Negative | Positive | Negative |
| | Chicken | 12 | Op | Negative | Negative | Negative | Negative |
| | | | C | Negative | Negative | Positive | Negative |
| | Chicken | 13 | Op | Negative | Negative | Negative | Negative |
| | | | C | Positive | Negative | Negative | Negative |
| | Chicken | 14 | Op | Negative | Negative | Negative | Negative |
| | | | C | Negative | Negative | Negative | Negative |
| | Chicken | 15 | Op | Positive | Positive | Positive | Negative |
| | | | C | Negative | Negative | Positive | Negative |
| | Chicken | 16 | Op | Positive | Positive | Positive | Negative |
| | | | C | Positive | Positive | Positive | Negative |
| | Chicken | 17 | Op | Positive | Positive | Positive | Negative |
| | | | C | Positive | Positive | Positive | Negative |
| | Chicken | 16,17 | Pooled Brain | Positive | Positive | Positive | Negative |
| Group B | Duck | 1 | Op | Positive | Positive | Positive | Negative |
| | | | C | Positive | Positive | Positive | Negative |
| | Duck | 2 | Op | Positive | Positive | Positive | Negative |
| | | | C | Negative | Negative | Positive | Negative |
| | Duck | 3 | Op | Positive | Positive | Positive | Negative |
| | | | C | Positive | Positive | Positive | Negative |
| | Duck | 4 | Op | Positive | Positive | Positive | Negative |
| | | | C | Positive | Positive | Positive | Negative |
| | Duck | 5 | Op | Positive | Positive | Positive | Negative |
| | | | C | Positive | Positive | Positive | Negative |
| | Duck | 6 | Op | Positive | Positive | Positive | Negative |
| | | | C | Positive | Positive | Positive | Negative |
| | Duck | 7 | Op | Positive | Positive | Positive | Negative |
| | | | C | Negative | Negative | Positive | Negative |
| | Duck | 8 | Op | Positive | Positive | Positive | Negative |
| | | | C | Negative | Negative | Positive | Negative |
| | Duck | 9 | Op | Positive | Positive | Positive | Negative |
| | | | C | Positive | Positive | Positive | Negative |
| | Duck | 18* | Op | Positive | Positive | Positive | Negative |
| | | | C | Positive | Positive | Positive | Negative |
| | Duck | 19* | Op | Positive | Positive | Positive | Negative |
| | | | C | Positive | Positive | Positive | Negative |
| | Goose | 20 | Op | Positive | Positive | Positive | Negative |
| | | | C | Positive | Positive | Positive | Negative |

*blood collected for serology, Duck 18 gave positive serology result (1/64) in HI using H5N1 clade 2.3.4.4b antigen

**Supplementary Table 4**: Serological assessment of sheep blood and milk samples taken from the infected premisses following detection of H5N1 HPAIV in poultry.

| **Collection date** | **Sheep identifier (sample type*)** | **HI** | **ELISA** | |
|---|---|---|---|---|
| | | **H5N1-clade 2.3.4.4b[#]** | **ID Vet High path H5** | **ID Vet Multispecies** |
| 7th March 2025 | Sheep 1 | <1/10 | Negative | Negative |
| | Sheep 2 | <1/10 | Negative | Negative |
| | Sheep 3 | <1/10 | Negative | Negative |
| | Sheep 4 | <1/10 | Negative | Negative |
| | Sheep 5 | <1/10 | Negative | Negative |
| | Sheep 6 | <1/10 | Negative | Negative |
| | Sheep 7 | **1/80** | **Positive** | **Positive** |
| | Sheep 8 | <1/10 | Negative | Negative |
| | Sheep 9 | <1/10 | Negative | Negative |
| | Sheep 10 | <1/10 | Negative | Negative |
| 14th March 2025 | Sheep 7 | **1/160** | **Positive** | **Positive** |
| | Sheep 7 (milk) | ND | **Positive** | **Positive** |
| 19th March 2025 | Sheep 7 | **1/80** | **Positive** | **Positive** |
| | Sheep 7 (milk) | ND | **Positive** | **Positive** |
| | Lamb 1[¥] | <1/10 | Negative | Negative |
| | Lamb 2[¥] | <1/10 | Negative | Negative |
| 21st March 2025 | Sheep 1 | <1/10 | ND | ND |
| | Sheep 2 | <1/10 | ND | ND |
| | Sheep 3 | <1/10 | ND | ND |
| | Sheep 4 | <1/10 | ND | ND |
| | Sheep 5 | <1/10 | ND | ND |
| | Sheep 6 | <1/10 | ND | ND |
| | Sheep 7 | <1/10 | ND | ND |
| | Sheep 8 | <1/10 | ND | ND |
| | Sheep 9 | <1/10 | ND | ND |
| | Sheep 10 | <1/10 | ND | ND |
| | Sheep 11 | <1/10 | ND | ND |
| | Sheep 12 | <1/10 | ND | ND |
| | Lamb 1 | <1/10 | ND | ND |
| | Lamb 2 | <1/10 | ND | ND |
| | Lamb 3 | <1/10 | ND | ND |
| | Lamb 4 | <1/10 | ND | ND |
| | Lamb 5 | <1/10 | ND | ND |
| | Lamb 6 | <1/10 | ND | ND |
| | Lamb 7 | <1/10 | ND | ND |
| 1st April 2025 | Sheep 1 | <1/10 | ND | ND |
| | Sheep 2 | <1/10 | ND | ND |
| | Sheep 3 | <1/10 | ND | ND |

| | | | | |
|---|---|---|---|---|
| | Sheep 4 | <1/10 | ND | ND |
| | Sheep 5 | <1/10 | ND | ND |
| | Sheep 6 | <1/10 | ND | ND |
| | Sheep 7 | <1/10 | ND | ND |
| | Sheep 8 | <1/10 | ND | ND |
| | Sheep 9 | <1/10 | ND | ND |
| | Sheep 10 | <1/10 | ND | ND |
| | Sheep 11 | <1/10 | ND | ND |
| | Sheep 12 | <1/10 | ND | ND |
| | Sheep 13 | <1/10 | ND | ND |
| | Sheep 14 | <1/10 | ND | ND |
| | Sheep 15 | <1/10 | ND | ND |
| | Sheep 16 | <1/10 | ND | ND |
| | Sheep 17 | <1/10 | ND | ND |
| | Sheep 18 | <1/10 | ND | ND |
| | Sheep 19 | <1/10 | ND | ND |
| | Sheep 20 | <1/10 | ND | ND |
| | Sheep 21 | <1/10 | ND | ND |
| | Sheep 22 | <1/10 | ND | ND |
| | Sheep 23 | <1/10 | ND | ND |
| | Sheep 24 | <1/10 | ND | ND |
| | Sheep 25 | <1/10 | ND | ND |

*Serum unless stated. [¥]both lambs were offspring of sheep #7. The Sheep sera were diluted due to RDE treatment. ND = test not performed. [#] A/chicken/Wales/053969/2021 (H5N1) antigen.

**Supplementary Table 5:** Summary of RT-PCR testing on sheep samples taken from the infected premisses following detection of H5N1 HPAIV in poultry.

| Sampling Date | Sample type | RT-PCR assay | | |
|---|---|---|---|---|
| | | M-gene | HP H5 | N1 |
| 7th March 2025 | Nasal swab | No Cq | No Cq | No Cq |
| | Rectal swab | No Cq | No Cq | No Cq |
| 14th March 2025 | Milk | **34.86** | **31.89** | **31.22** |
| | Nasal swab | No Cq | No Cq | No Cq |
| | Rectal swab | No Cq | No Cq | No Cq |
| 19th March 2025 Swab and liquid samples PM | Milk | **34.02** | **33.55** | **32.98** |
| | Blood (EDTA) | No Cq | No Cq | No Cq |
| | Nasal swab | No Cq | No Cq | No Cq |
| | Rectal swab | No Cq | No Cq | No Cq |
| | External teat left swab | No Cq | No Cq | No Cq |
| | External teat right swab | No Cq | No Cq | No Cq |
| | Intra teat left swab | No Cq | No Cq | No Cq |
| | Intra teat right swab | No Cq | No Cq | No Cq |
| | Pharyngeal swab | No Cq | No Cq | No Cq |
| | Skin & wool swab | No Cq | No Cq | No Cq |
| | Tracheal swab 1 | No Cq | No Cq | No Cq |
| | Tracheal swab 2 | No Cq | No Cq | No Cq |
| 19th March 2025 Tissue samples PM | Abomasum | No Cq | No Cq | No Cq |
| | Bladder | No Cq | No Cq | No Cq |
| | Brain | No Cq | No Cq | No Cq |
| | Bronchial section | No Cq | No Cq | No Cq |
| | Colon | No Cq | No Cq | No Cq |
| | Duodenum | No Cq | No Cq | No Cq |
| | Gall Bladder | No Cq | No Cq | No Cq |
| | Heart | No Cq | No Cq | No Cq |
| | Ileum | No Cq | No Cq | No Cq |
| | Jejunum | No Cq | No Cq | No Cq |
| | Kidney | No Cq | No Cq | No Cq |
| | Liver | No Cq | No Cq | No Cq |
| | Lung | No Cq | No Cq | No Cq |
| | Lymph nodes Pool | No Cq | No Cq | No Cq |
| | Mammary gland -Left | No Cq | No Cq | No Cq |
| | Mammary gland -Right | No Cq | No Cq | No Cq |
| | Oesophagus | No Cq | No Cq | No Cq |
| | Omasum | No Cq | No Cq | No Cq |
| | Ovary | No Cq | No Cq | No Cq |
| | Pancreas | No Cq | No Cq | No Cq |
| | Pharyngeal Mucosa | No Cq | No Cq | No Cq |
| | Reticulum | No Cq | No Cq | No Cq |
| | Rumen | No Cq | No Cq | No Cq |
| | Spleen | No Cq | No Cq | No Cq |
| | Tracheal pool 1 | No Cq | No Cq | No Cq |
| | Tracheal pool 2 | No Cq | No Cq | No Cq |
| | Turbinate | No Cq | No Cq | No Cq |
| | Uterus | No Cq | No Cq | No Cq |

| | | | | |
|---|---|---|---|---|
| 19th March 2025 Swab samples from lambs | Lamb 1- Oral Swab | No Cq | No Cq | No Cq |
| | Lamb 1- Rectal Swab | No Cq | No Cq | No Cq |
| | Lamb 2- Oral Swab | No Cq | No Cq | No Cq |
| | Lamb 2- Rectal Swab | No Cq | No Cq | No Cq |
| 21St March swab samples from sheep and lambs | Sheep 1-12 - Nasal swab | No Cq | No Cq | No Cq |
| | Sheep 1-12 - Rectal swab | No Cq | No Cq | No Cq |
| | Sheep 4,7,10 - Milk | No Cq | No Cq | No Cq |
| | Lamb 1-7- Nasal swab | No Cq | No Cq | No Cq |
| | Lamb 1-7- Rectal swab | No Cq | No Cq | No Cq |
| 1st April 2025 Swab samples from remaining sheep and lambs | Sheep 1-25 Nasal | No Cq | No Cq | No Cq |
| | Sheep 1-25 Rectal | No Cq | No Cq | No Cq |
| | Sheep 1,2,4 Milk Left teat | No Cq | No Cq | No Cq |
| | Sheep 1,2,3,4 Milk Right teat | No Cq | No Cq | No Cq |

**Supplementary Table 6:** Amino acid differences detected in the viral RNA detected in the sheep compared to the sequence derived from the birds on the infected premises.

| Segment | Amino acid position | Avian sequences* | Sheep #7 (milk) |
|---|---|---|---|
| PB2 | 456 | N | D |
| PA | 335 | L | F |
| NA | 75 | L | F |
| | 114 | V | M |
| HA | 171[a] (155[b] / 159[c]) | D | N |
| | 277[a] (261[a] / 264[b]) | D | G |
| PB1 | 290 | M | V |
| | 577 | K | E |
| | 688 | Q | H |

*Compared to all avian sequences derived from the infected premises; [a] immature H5 HA numbering; [b] mature H5 HA numbering; [c] mature H3 numbering.
